# Invasion performance-similarity found among multiple cell systems

**DOI:** 10.1101/2020.08.15.252445

**Authors:** Om Prakash

## Abstract

Understanding of inter-system behavior develops biologically relevant intuition for drug repositioning as well as other biological research. But combining all the possible genes interactions into a system, and furthermore comparisons of multiple systems are a challenge on time ground with feasible experiments. In present study, 64 cell lines from 11 different organs were compared for their invasion performance. RNA expressions of 23 genes were used to create systems artificial neural network (ANN) models. ANN models were prepared for all 64 cell lines and observed for their invasion performance through network mapping. The resulted cell line clusters bear feasible capacity to perform experiments for biologically relevant research motivations as drug repositioning and selective targeting etc.; and can be used for analysis of invasion related aspects.

## INTRODUCTION

Perceptive for inter-system behavior is useful for drug repositioning, selective targeting as well as other biological research. Although cellular components (i.e. genes) are same in different tissues; but systems function differently in different tissues. Combining all the possible genes interactions into a system, and furthermore comparisons of multiple systems are a challenge on time ground with feasible experimentally.

Invasion property has been selected to understand the systems similarity for specific function of cell. Cell invasion is the property linked with its migration, and making it able to be motile through extracellular matrix. Cancer cells become invasive and disseminate to tissue to make metastases. Multiple gene combinations have been found to be involved for cell invasion.

Some of the gene expressions known in relation with cell invasion are as: Role of T cell lymphoma invasion and metastasis 1 (TIAM1) was observed on tumor cell invasion and metastases (Xu et. al. 2010). KiSS-1 metastasis suppressor (KISS1) was found as mediator in triple negative breast cancer (Tian et. al. 2018). Anti S100 calcium binding protein A4 (S100A4) was found to suppress cell invasion (Klingelhöfer et. al. 2012). Cortactin (CTTN) was observed for promotion of cell invasion in colorectal cancer (Jing et. al. 2016). Protein arginine methyltransferase 7 (PRMT7) is known for metastasis in human non-small-cell lung cancer (Cheng et. al. 2018). Matrix metallopeptidase 26 (MMP26) was found to regulate chondrosarcoma (Xu et. al. 2015). Epithelial stromal interaction 1 (EPSTI1) is reported for breast cancer invasion and metastasis (Tianshu et. al. 2014). Neural precursor cell expressed, developmentally down-regulated 9 (NEDD9) was found in relation with cell invasion in colon cancer (Jin et. al. 2016). CUB domain containing protein 1 (CDCP1) regulates invasion in cancer cells (Miyazawa et. al. 2013). SEC11 homolog A, signal peptidase complex subunit (SEC11A) was found to linked with gastric cancer via TGF-alpha (Oue et. al. 2014). Reversion inducing cysteine rich protein with kazal motifs (RECK) inhibits cervical cancer through p53 pathway (Liu et. al. 2018). EPH receptor B6 (EPHB6) was found to alter cell invasion in breast cancer (Fox et. al. 2009).

Application of artificial neural network (ANN) modeling is well established in pharmaceutical and medical research. It has shown great potential to classify large biological data (Khan et. al. 1998; Agatonovic-Kustrin et. al. 2000; Hoffman et. al 2013). Therefore in present study, invasion performance has been clustered among multiple cell systems. It is an ANN implemented network mapping of invasion related RNA expressions in 64 systems from 11 different organs.

## MATERIALS & METHODS

### Initial dataset

System’s RNA expressions of multiple genes were collected for multiple cell lines. RNA expression were collected for multiple tissue systems (64 cell lines from 11 different organ systems) as: Abdominal (cell lines: CACO-2, CAPAN2, HepG2), Brain (cell lines: AF22, SH-SY5Y, U-138 MG, U-251 MG, U-87 MG), Endothelial (cell lines: hTEC/SVTERT24-B, HUVEC TERT2, TIME), Female Reproductive organ systems (cell lines: AN3-CA, EFO-21, HeLa, hTERT-HME1, MCF7, SiHa, SK-BR-3, T-47d), Fibroblast (cell lines: BJ, BJ hTERT+, BJ hTERT+ SV40 Large T+, BJ hTERT+ SV40 Large T+ RasG12V, fHDF/TERT166, HBF TERT88), Lung (A549, HBEC3-KT, SCLC-21H), Lymphoid (cell lines: Daudi, HDLM-2, Karpas-707, MOLT-4, REH, RPMI-8226, U-266/70, U-266/84, U-698), Myeloid (cell lines: HAP1, HEL, HL-60, HMC-1, K-562, NB-4, THP-1, U-937), Renal (cell lines: HEK 293, NTERA-2, PC-3, RPTEC TERT1, RT4), Sarcoma (cell lines: ASC diff, ASC TERT1, LHCN-M2, RH-30, U-2 OS, U-2197), Skin (cell lines: A-431, HaCaT, SK-MEL-30, WM-115), and some Miscellaneous (cell lines: BEWO, HHSteC, HSkMC, hTCEpi). Collections were made from Human Protein Atlas.

Gene expressions used for systems mapping for invasion performance were as: T cell lymphoma invasion and metastasis 1 (TIAM1), KiSS-1 metastasis suppressor (KISS1), S100 calcium binding protein A4 (S100A4), Cortactin (CTTN), Twist family bHLH transcription factor 1 (TWIST1), T cell lymphoma invasion and metastasis 2 (TIAM2), Sperm adhesion molecule 1 (SPAM1), S100 calcium binding protein A11 (S100A11), CD151 molecule (Raph blood group) (CD151), Cadherin 1 (CDH1), Suppression of tumorigenicity 14 (ST14), Kallikrein related peptidase 7 (KLK7), Cadherin 20 (CDH20), Matrix metallopeptidase 25 (MMP25), Protein arginine methyltransferase 7 (PRMT7), Matrix metallopeptidase 26 (MMP26), Epithelial stromal interaction 1 (EPSTI1), Neural precursor cell expressed, developmentally down-regulated 9 (NEDD9), CUB domain containing protein 1 (CDCP1), SEC11 homolog A, signal peptidase complex subunit (SEC11A), Reversion inducing cysteine rich protein with kazal motifs (RECK), EPH receptor B6 (EPHB6), and Tetratricopeptide repeat domain 9 (TTC9).

### Method used for ANN based mapping of systems gene expressions

Since no initial classification criterion was available for invasion performance of cell lines, therefore a customized training of ANN model was adopted. ANN model was supervised with ‘Bipolar Rotational Targets’. A detail about algorithm is as following:

***Customized training of Artificial Neural Network:*** ANN architecture of ‘Input-Hidden-Output’ was implemented for model development. Although ANN architecture was supervised; but customized training was processed with ‘bipolar rotational target’ (BRT) theme. BRT was introduced as that each sample was trained with both binary values ‘0’ and ‘1’, but in rotational manner for multiple times. Multiple models were prepared with multiple seed values.
***Post-training processing:*** Multi-model outputs were processed to calculate cluster-score for each sample. Cluster-score is a z-score values identified through nested processing of Multi-model outputs. Cluster score has been defined as:

Let’s Sample set: {S_1 to M_}_N_

Where ‘N’ is number of samples (S); and each sample has ‘M’ number of outputs from ‘M’ ANN-models. For each sample vector; mean {μ_N_} and standard deviation {σ_N_} were calculated. Furthermore mean (μ’) and standard deviation (σ’) was calculated for {(μ/σ)_N_}

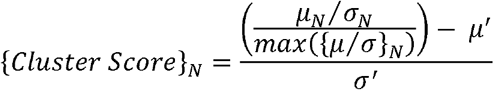
***Network Graph with node values as cluster-score*** Cluster-score used as target tags for network mapping. Python package ‘Networkx’ was used for network graph preparation and analysis. Since ANN model capture core relation between data points, therefore it was supposed to get the highest linked interactions into zone of least distant nodes; and low linked interactions in zone of maximal distant nodes.

## RESULTS AND DISCUSSION

Combined Network graph of multiple systems (11 organs and 64 cell lines) prepared for observation of relative invasion performance-similarity. All-node combination network graph was filtered for maximum confidant edge compactness with distance of 0.02 only. Five interlinked clusters were observed (Figure 1 a, b). For cross evaluation, 03 separate clusters for three different systems were plotted in Figure 2. Figure 2(a) showed that, cluster centered for invasion performance of Lung cell line A549, at node distance of 0.02. Lung system was found to be clustered with Abdominal, Lymphoid, Brain, Fibroblast, Renal, Myeloid, Sarcoma and cell line from Female reproductive organ. Experimental evidences found for explanation of A549 and Caco-2 in relation of cell adhesion (Cheng et. al. 2015). Figure 2(b) showed that, cluster centered for invasion performance of Brain cell line U-87 MG, at node distance of 0.02. Brain system was found to be clustered with Fibroblast, Myeloid, Lung and Lymphoid systems. Brain cell expressions have been found to affect lymphoid and female reproductive organs. Since, biology of malignancy of cancer cells is directly related with cell adhesion molecules and transition of clustered epithelial cells into free-to-move mesenchymal cells. Two genes NLGN4X (for neuronal adhesion) and CD66 (for cervical cancers) are known for their effect on malignancy in epithelial cancer cells. In some studies, impact of NLGN4X was also observed on lymphoid tissue. Epithelial to mesenchymal transitions are related with prolyl 4-hydroxylase 9 (P4H9) and glycosylation at molecular level. Neuroligin is known to affect lymphoid cells. Figure 2(c) showed that, cluster centered for invasion performance of female reproductive organ cell line MCF7, at node distance of 0.02. The system was found to be clustered with Skin, Abdominal and Sarcoma cell lines. Experimental evidences found for Neuroligin affects metastatic performance of CD66+ cells (doi: https://doi.org/10.1101/2020.07.29.226274), which include many cell lines from female reproductive organs.

**Figure 1(a).**
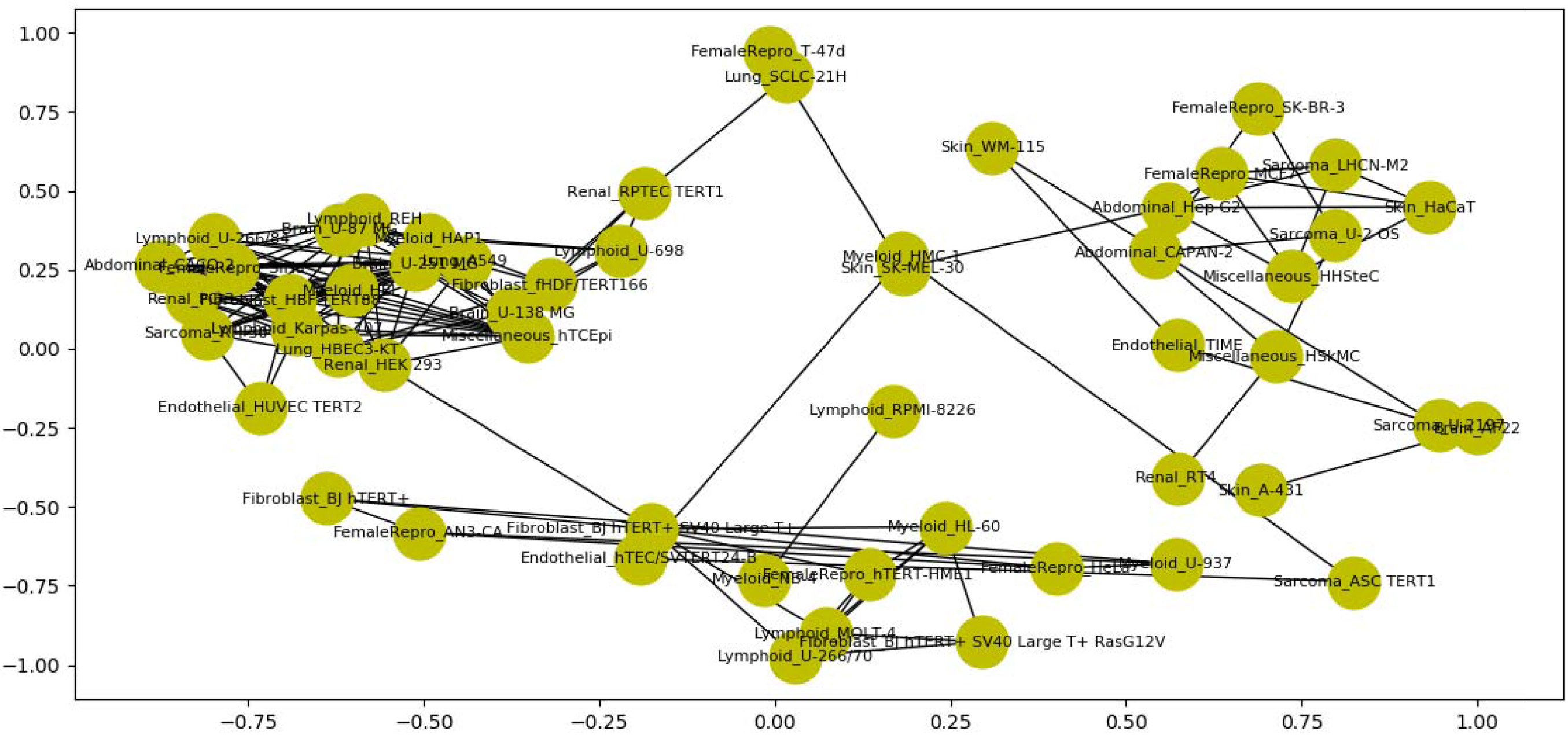
Combined Network graph of multiple systems (11 organs & 64 cell lines) prepared for observation of relative invasion performance-similarity. Graph was filtered for maximum confidant edge compactness with distance of 0.02 only.

**Figure 1(b).**
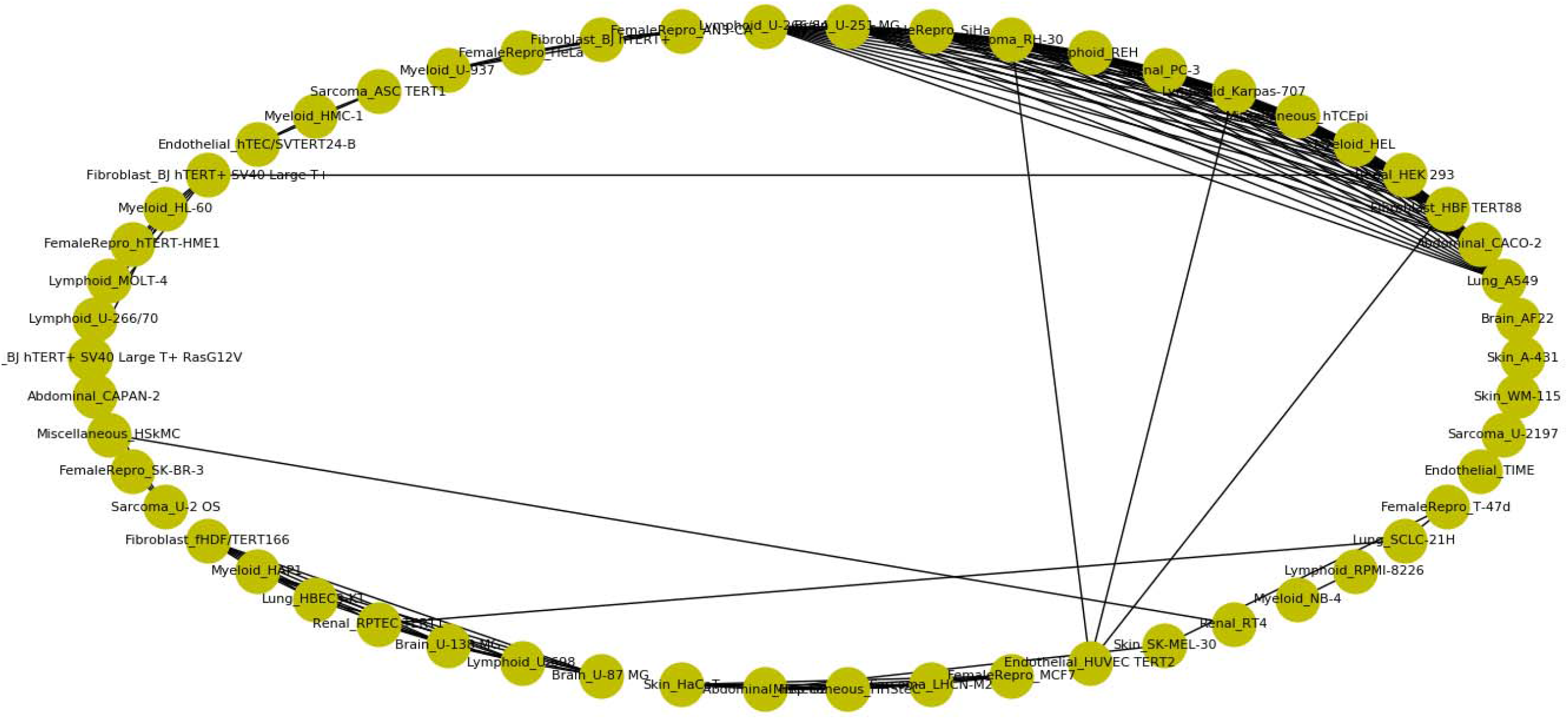
Circular Network graph of multiple systems (11 organs & 64 cell lines) prepared for observation of relative invasion performance-similarity. Graph was filtered for maximum confidant edge compactness with distance of 0.02 only.

**Figure 2(a).**
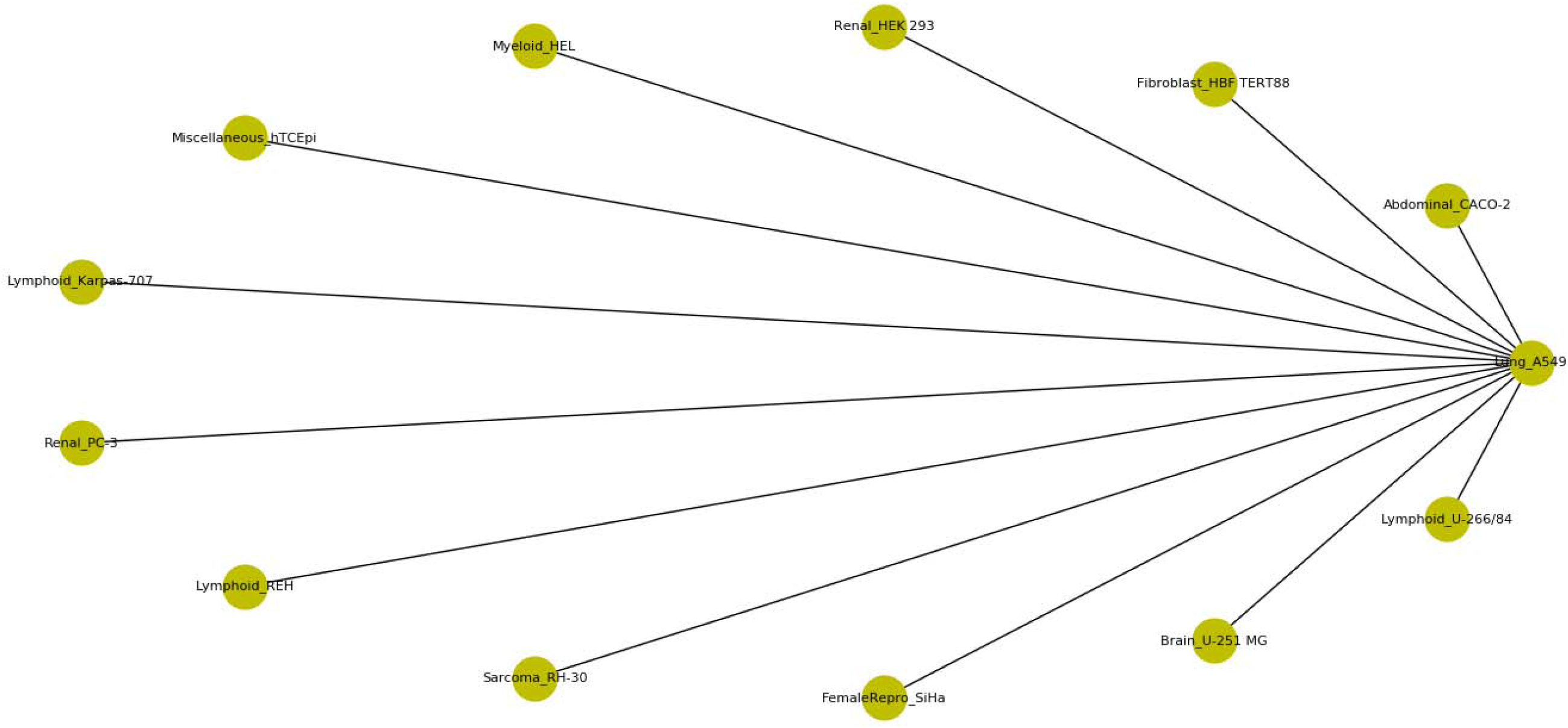
Cluster centered for invasion performance of Lung cell line A549, at node distance of 0.02. Lung system was found to be clustered with Abdominal, Lymphoid, Brain, Fibroblast, Renal, Myeloid, Sarcoma and cell line from Female reproductive organ.

**Figure 2(b).**
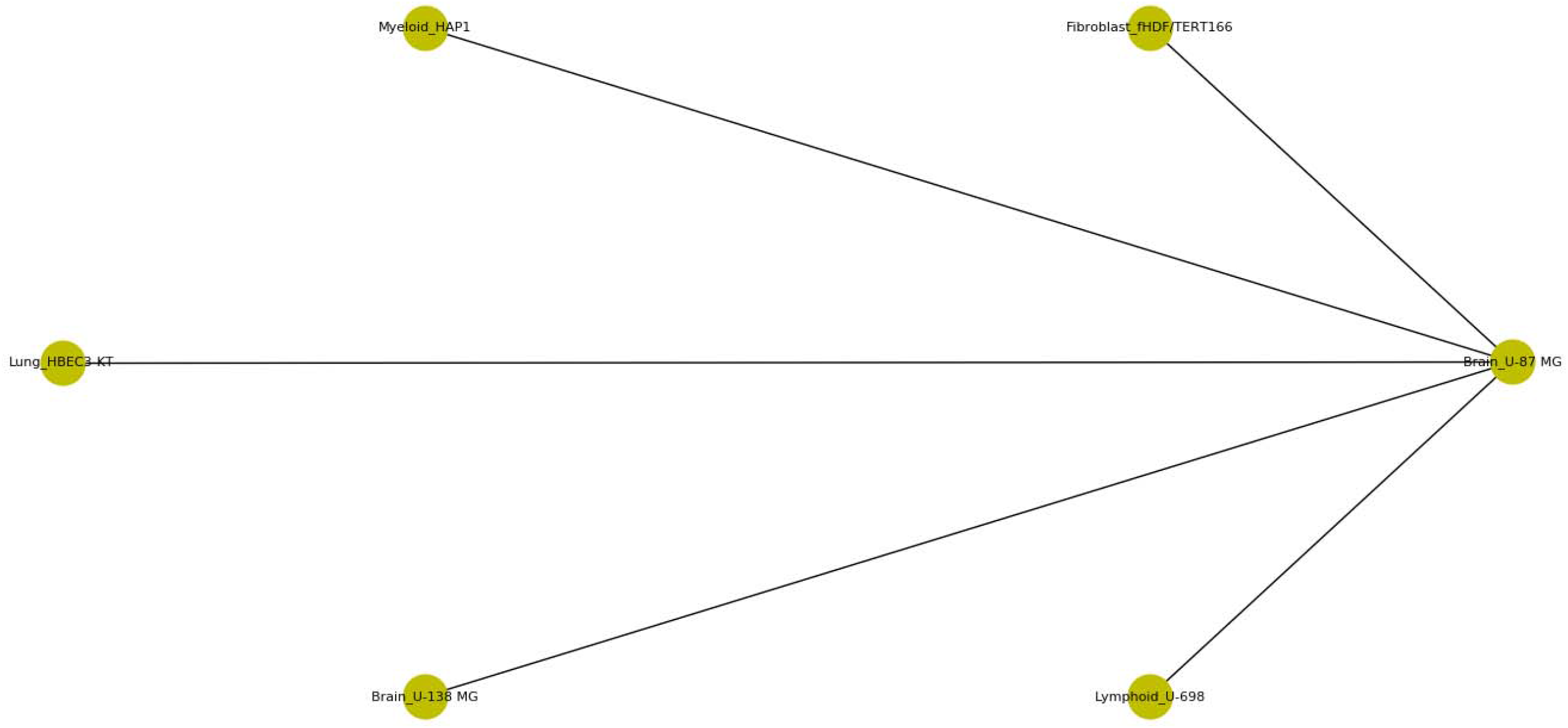
Cluster centered for invasion performance of Brain cell line U-87 MG, at node distance of 0.02. Brain system was found to be clustered with Fibroblast, Myeloid, Lung and Lymphoid systems.

**Figure 2(c).**
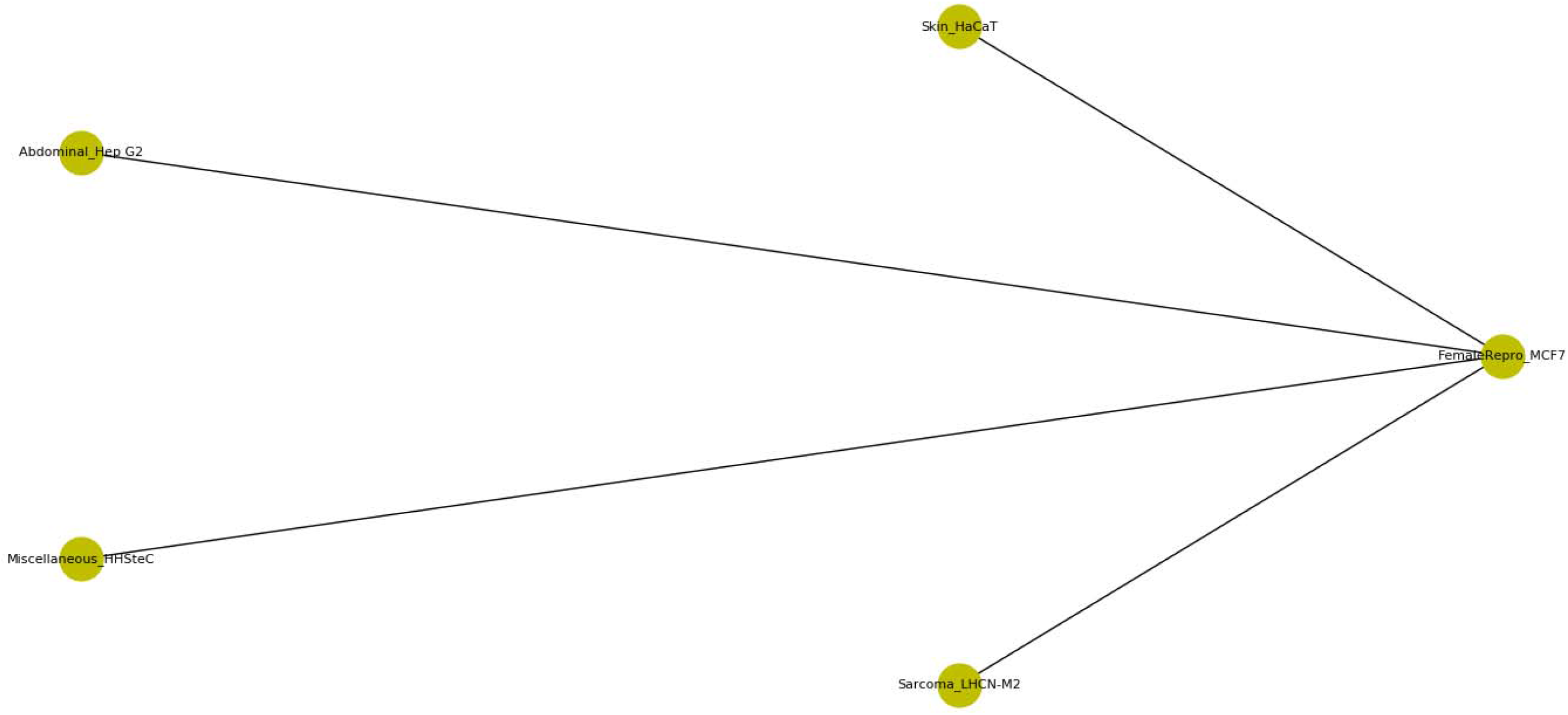
Cluster centered for invasion performance of Female reproductive organ cell line MCF7, at node distance of 0.02. The system was found to be clustered with Skin, Abdominal and Sarcoma cell lines.

## CONCLUSION

Invasion performance of multiple cell systems (64 systems from 11 organs) were clustered through Artificial Neural Network based systems models. Indications were found about possibilities of existence of similar mechanism into different tissue cells. These indications will be benefitted as supportive knowledge for drug repositioning, selective targeting and other R&Ds in molecular biology.

## ACKNOWLEDGEMENT

Author express gratitude to *The Institute of Mathematical Sciences*, Chennai-600113, India for providing research facilities as well as DAE Post-Doctoral Fellowship (PDF 214).

## REFERENCES

Xu, K., Rajagopal, S., Klebba, I. et al. The role of fibroblast Tiam1 in tumor cell invasion and metastasis. Oncogene 29, 6533–6542 (2010). https://doi.org/10.1038/onc.2010.385.

Tian J, Al-Odaini AA, Wang Y, et al. KiSS1 gene as a novel mediator of TGFβ-mediated cell invasion in triple negative breast cancer. Cell Signal. 2018;42:1–10. doi:10.1016/j.cellsig.2017.10.002.

Jörg Klingelhöfer, Birgitte Grum-Schwensen, Mette K Beck, Rikke Stagaard Petersen Knudsen, Mariam Grigorian, Eugene Lukanidin, Noona Ambartsumian. Anti-S100A4 Antibody Suppresses Metastasis Formation by Blocking Stroma Cell Invasion. Neoplasia. 2012 Dec; 14(12): 1260–1268. doi: 10.1593/neo.121554.

Jing X, Wu H, Ji X, Wu H, Shi M, Zhao R. Cortactin promotes cell migration and invasion through upregulation of the dedicator of cytokinesis 1 expression in human colorectal cancer. Oncol Rep. 2016;36(4):1946–1952. doi:10.3892/or.2016.5058

Cheng D, He Z, Zheng L, Xie D, Dong S, Zhang P. PRMT7 contributes to the metastasis phenotype in human non-small-cell lung cancer cells possibly through the interaction with HSPA5 and EEF2. Onco Targets Ther. 2018;11:4869–4876. Published 2018 Aug 14. doi:10.2147/OTT.S166412.

Xu X, Ma J, Li C, Zhao W, Xu Y. Regulation of chondrosarcoma invasion by MMP26. Tumour Biol. 2015;36(1):365–369. doi:10.1007/s13277-014-2657-7.

Tianshu Li, Heng Lu, Chao Shen, Satadru K. Lahiri, Melissa S. Wason, Debarati Mukherjee, Lin Yu, and Jihe Zhao. Epithelial stromal interaction 1 (EPSTI1) is reported for breast cancer invasion and metastasis: Identification of epithelial stromal interaction 1 as a novel effector downstream of Krüppel-like factor 8 in breast cancer invasion and metastasis.. Oncogene. 2014 Sep 25; 33(39): 4746–4755.

Jin Dai, Peter G. Van Wie, Leonard Yenwong Fai, Donghern Kim, Lei Wang, Pratheeshkumar Poyil, Jia Luo, and Zhuo Zhanga,. Downregulation of NEDD9 by apigenin suppresses migration, invasion, and metastasis of colorectal cancer cells.Toxicol Appl Pharmacol. 2016 Nov 15; 311: 106–112.

Y, Uekita T, Ito Y, Seiki M, Yamaguchi H, Sakai R. CDCP1 regulates the function of MT1-MMP and invadopodia-mediated invasion of cancer cells. Mol Cancer Res. 2013;11(6):628–637. doi:10.1158/1541-7786.MCR-12-0544.

Oue, N., Naito, Y., Hayashi, T. et al. Signal peptidase complex 18, encoded by SEC11A, contributes to progression via TGF-α secretion in gastric cancer. Oncogene 33, 3918–3926 (2014). https://doi.org/10.1038/onc.2013.364.

Liu Y, Li L, Liu Y, et al. RECK inhibits cervical cancer cell migration and invasion by promoting p53 signaling pathway. J Cell Biochem. 2018;119(4):3058–3066. doi:10.1002/jcb.26441.

Fox, B., Kandpal, R. EphB6 receptor significantly alters invasiveness and other phenotypic characteristics of human breast carcinoma cells. Oncogene 28, 1706–1713 (2009). https://doi.org/10.1038/onc.2009.18.

Agatonovic-Kustrin S, Beresford R. Basic concepts of artificial neural network (ANN) modeling and its application in pharmaceutical research. J Pharm Biomed Anal. 2000;22(5):717–727. doi:10.1016/s0731-7085(99)00272-1.

Khan ZH, Mohapatra SK, Khodiar PK, Ragu Kumar SN. Artificial neural network and medicine. Indian J Physiol Pharmacol. 1998;42(3):321–342.

Hoffman MR, Mielens JD, Omari TI, Rommel N, Jiang JJ, McCulloch TM. Artificial neural network classification of pharyngeal high-resolution manometry with impedance data. Laryngoscope. 2013;123(3):713–720. doi:10.1002/lary.23655.

Cheng XD, Gu JF, Yuan JR, Feng L, Jia XB. Suppression of A549 cell proliferation and metastasis by calycosin via inhibition of the PKC-α/ERK1/2 pathway: An in vitro investigation [published correction appears in Mol Med Rep. 2016 Apr;13(4):3709–10]. Mol Med Rep. 2015;12(6):7992-8002. doi:10.3892/mmr.2015.4449.

Om Prakash. Neuroligin affects metastatic performance of CD66+ cells: Theoretical analysis of Multi-tissue RNA expressions through System modeling based Network mapping under in-vitro constraints. bioRxiv 2020.07.29.226274; doi: https://doi.org/10.1101/2020.07.29.226274.

